# Deep Spatial Sequencing Revealing Differential Immune Responses in Human Hepatocellular Carcinoma

**DOI:** 10.1101/2025.03.25.645292

**Authors:** Yan-Ping Yu, Silvia Liu, Caroline Obert, Bao-Guo Ren, Marielle Krivet, Kyle Metcalfe, Jia-Jun Liu, Tuval Ben-Yehezkel, Jian-Hua Luo

## Abstract

Hepatocellular carcinoma (HCC) is one of the most lethal cancers for humans. HCC is highly heterogeneous. In this study, we performed ultra-depth (∼1 million reads per spot) spatial sequencing on a case of HCC. Sixteen distinct spatial expression clusters were identified. Each of these clusters was spatially contiguous and had distinct gene expression patterns. In contrast, benign liver tissues showed minimal heterogeneity in terms of gene expression. Numerous immune cell-enriched spots were identified in both HCC and benign liver regions. Cells adjacent to these immune cell-enriched spots showed significant alterations in their gene expression patterns. Interestingly, the responses of HCC cells to the nearby immune cells were significantly more intense and broader, while the responses of benign liver cells to immune cells were somewhat narrow and muted, suggesting an innate difference in immune cell activities towards HCC cells in comparison with benign liver cells. When standard-depth sequencing was performed, significant numbers of genes and pathways that were associated with these changes disappeared. Qualitative differences in some pathways were also found. These results suggest that deep spatial sequencing may help to uncover previously unidentified mechanisms of liver cancer development.

## Introduction

Liver cancer is one of the most lethal malignancies for humans and causes over 700,000 deaths worldwide annually^1-3^. Hepatocellular carcinoma (HCC) is the most common type of liver cancer and accounts for 90% of all liver cancers^4,5^. Large numbers of genomic and gene expression abnormalities have been discovered in HCC^6-19^. HCC has high levels of heterogeneity. These heterogeneities may result from the underlying variation of genomic alterations. However, the location-based heterogeneity of HCC has rarely been analyzed at the genetic level.

In recent years, spatial genetic analysis has been rapidly developed. Cancer microenvironment has been shown to have significant variations in different cancer locations^20^. Studies utilizing spatial genetic analysis revealed spatial relationships of cell-cell interaction and the impact of genetic alterations of a cell on its adjacent tissue microenvironment^21^. Due to the significant genetic heterogeneity in an HCC sample, the tumor microenvironment may differ from region to region. In this report, we performed a 10x Genomics Visium Spatial transcriptome analysis on a case of HCC samples. Significant variation of gene expression patterns was found in different regions of the cancer samples.

## Materials and Methods

### Tissue samples and CytAssist workflow

A case of HCC sample with moderate differentiation and a history of alcohol abuse and Hepatitis B infection was obtained from archived tissue slide storage. The case was fully anonymized. The tissue procurement protocols were approved by the Institutional Review Board of University of Pittsburgh. All procedure and protocols were carried out in accordance with the guidelines by the Institutional Review Board of University of Pittsburgh. The cancer and benign liver regions were identified by a board-certified pathologist. The slides were deparaffinized, Hematoxylin/Eosin stained, and underwent Visium cassette assembly, probe hybridization/ligation, tissue transfer to Visium slide, DNA isolation, clean-up, and ligation probe amplification based on the manufacturer’s manual. Standard sequencing was performed on Illumina NextSeq 550 Dx platform, while ultra-depth sequencing was performed on the Element Biosciences AVITI platform. The sequencing procedures followed the manufacturers’ recommendations^22,23^.

### Statistical analysis for Spatial transcriptomics data

Spatial transcriptomics was measured by the Visium CytAssist platform. The H&E staining file was imported to the Loupe Browser (10x Genomics) for spatial alignment. Then, the alignment file and the raw sequencing FASTQ files were processed by Space Ranger (10x Genomics) to align to the human reference genome hg38. After pre-processing, the feature by cell count matrix and the imaging files were analyzed by the R Seurat package^24^. To integrate two slides, top 3000 high-variable genes to integrate the two libraries using SCT transformation. Principal component analysis followed by the Uniform Manifold Approximation and Projection (UMAP) was applied for dimension reduction and visualization. Spatial spots were clustered based on the gene expression profiles. Spatial transcriptomics data were visualized with UMAP and spatial feature plot provided by the Seurat package.

To check the immune cells and their tissue microenvironments, spots with high immune expression were identified and further analyzed. Kupffer cells were identified by CD68, CD163, LYZ, C1QA, AIF1; T cells were identified by CD3D, CD2, IL7R, TRBC2, CD69; B cells and Plasma cells were defined by IGKC, JCHAIN, CD79A, CD27, CD74; NK and other immune cells were defined by CD4, CD8A, ITGAM, NKG7, KLRD1, PRF1, CD7, TRDC. The immune cell-enriched spots were defined by the average expression of the above immune markers higher than 0.9. The spots adjacent to immune cells were defined as the spots in contact with the immune cells. All the other spots were labeled as non-immune spots. In addition, HCC and benign liver spots were identified based on morphology in the H&E staining. Differential expression analyses were performed comparing (1) HCC adjacent to versus away from immune cell-enriched spots; (2) Benign liver cells adjacent to versus away from immune cell-enriched spots; (3) HCC cells adjacent to immune spots versus benign liver cells adjacent to immune spots. Further, top genes were screened by integrating these differentially expressed genes with the top 3000 high-variable genes. These genes were then used for Ingenuity Pathway Analysis.

The spatial transcriptomics data were further integrated with the public single-cell RNA-seq data^25^, which contains samples from the HCC tumor cores, tumor borders, and adjacent non-tumor tissues. Spatial deconvolution was performed and visualized by tool CARD ^26^ to spatially infer the proportions of different immune cell types per immune spot.

## Data Availability

The spatial transcriptomics data were submitted to the Gene Expression Omnibus (GEO) database with accession ID GSE283406.

## Results

To analyze the spatial transcriptomes and gene expression alteration patterns in a space-related fashion, we selected a case of moderately differentiated HCC containing multiple cancer nodules. As shown in Figure 1A, two slides were analyzed for the spatial gene expression. One of the slides contained some benign liver tissues adjacent to the cancer, while the other slide contained only liver cancer tissues. Deep sequencing was performed to reach 697,228 to 1,327,551 mean reads per spot, detecting 1465 to 3223 median genes per spot. Two slides account for 6320 spots of tissues. A Seurat package was employed to identify marker genes that clustered the spots. Sixteen distinct clusters of spots were identified based on the top 3000 highly variable gene expressions (Figure 1B-D, supplemental table 1). The distributions of these spot clusters on the slides did not appear random but rather aggregated in a distinct, patchy manner. Since most of the spots were cancer cells dominated. These clusters may reflect gene expression variations among the cancer cells. Pathway analyses (supplemental tables 2-17) showed that the areas of benign liver tissues were dominated by gene expression of biosynthesis of cholesterol (supplemental table 11). They were mostly aggregated as a cluster (cluster 9, figure 1B and D). On the other hand, high heterogeneity for the areas of HCC was identified: Fifteen distinctive clusters were found (Figures 1C and D). Some have characteristics of fibrotic gene expression patterns (clusters 0, 6, 14, and 15, supplemental tables 2, 8, 16, and 17), while the others were also dominated by matrix/cell-cell contact activation pathways (clusters 3, 5, 8, 14, and 15, supplemental tables 5, 7, 10, 16, and 17). Pro-growth pathways were found to overexpress in clusters 1, 3, 8, and 11(Supplemental tables 3, 5, 10, and 13). Genes responsible for coagulation activation, one of the key features of HCC, were found overexpressed in clusters 2, 10, and 11 (Supplemental tables 4, 12, and 13). Many of the clusters overexpressed genes essential for oxidative metabolism (clusters 4, 5, 6, 7, 10, 12, and 13, supplemental tables 6-9, 12, 14, and 15). Their spatial distribution in the slides was aggregated in a contiguous patchy fashion, suggesting that they may arise as clonal expansions from single cells.

**Figure 1.**
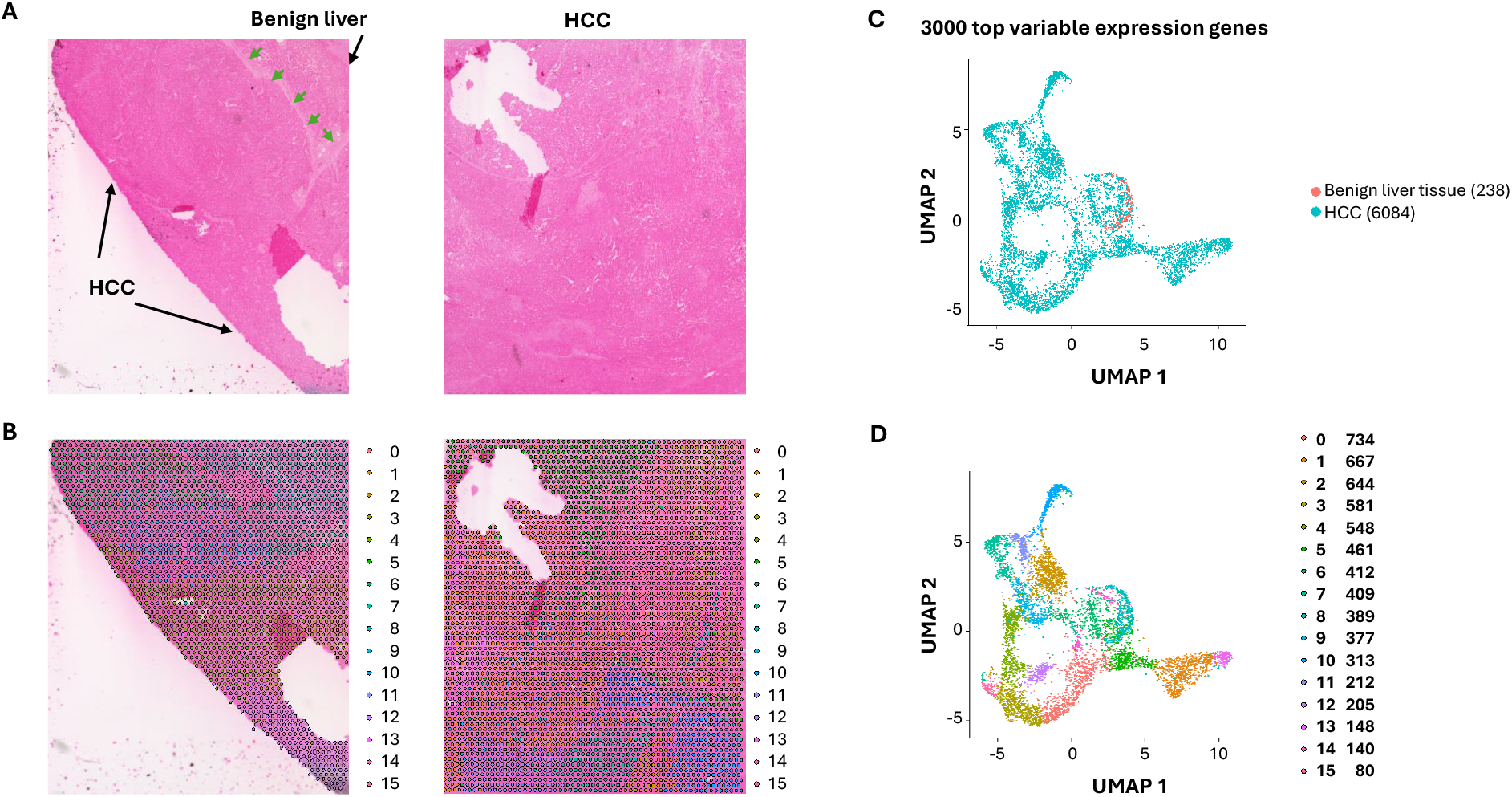
Spatial distribution of HCC clones with ultra-depth sequencing. (A) Hematoxylin and Eosin staining of HCC and its adjacent benign tissues. The areas of benign liver tissues and HCC are indicated. The demarcation between the benign liver and HCC is indicated by arrows. (B) The UMAP distribution of benign liver and HCC cells. The HCC and benign liver cell clusters are indicated. (C) Spatial visualization of cell clusters in slides. Each cluster distribution is indicated by its unique color. (D) UMAP distributions of 16 spatial clusters.

### Identification of immune cell-enriched spots

The cancer microenvironment plays a critical role in shaping cancer development. Significant lymphocytes and macrophages were identified in both benign liver cancer and HCC areas. To investigate the impact of these immune cells on liver cancer, selected gene markers for cells of immune lineages including Kuffler cells (Cd68, Cd163, Lyz, C1qa, Aif1), T cells (Cd3d, Cd2, Il7r, Trbc2, Cd69, Cd4, Cd8, Cd11), B cells (Igkc, Jchain, Cd79a, Cd27, Cd74) and NK (Nkg7, Klrd1, Prf1, Cd7, Trdc) cells were analyzed for each spot. Spots with average expression of these markers above 0.9 were deemed immune cell-enriched spots. Two hundred eighty-nine spots were identified as immune cell-enriched. As shown in Figure 2A, many immune cell-enriched spots were located in the benign liver tissues adjacent to HCC, while the distribution of immune cell-enriched spots in the HCC area varied from region to region. B cells and myeloid cells were the predominant cell types in the immune cell-enriched spots in both HCC and benign liver areas (Supplemental Figure 1). Interestingly, all types of immune cells were more abundant in the immune cell-enriched spots in the benign liver (Supplemental Figure 2). Most immune cell-enriched spots co-localized with clusters 3 and 9 (figure 2B). Next, we categorized the immune microenvironment into cells that were adjacent to immune cell-enriched spots (adjacent to immune cells) versus those that were away from the immune cells (away from immune cells) (Figures 2A and B). Six hundred seventy spots were deemed impacted by location adjacent to immune cells, including 84 spots from benign liver and 586 from HCC. Spots away from immune cell-enriched spots were 5361.

**Figure 2.**
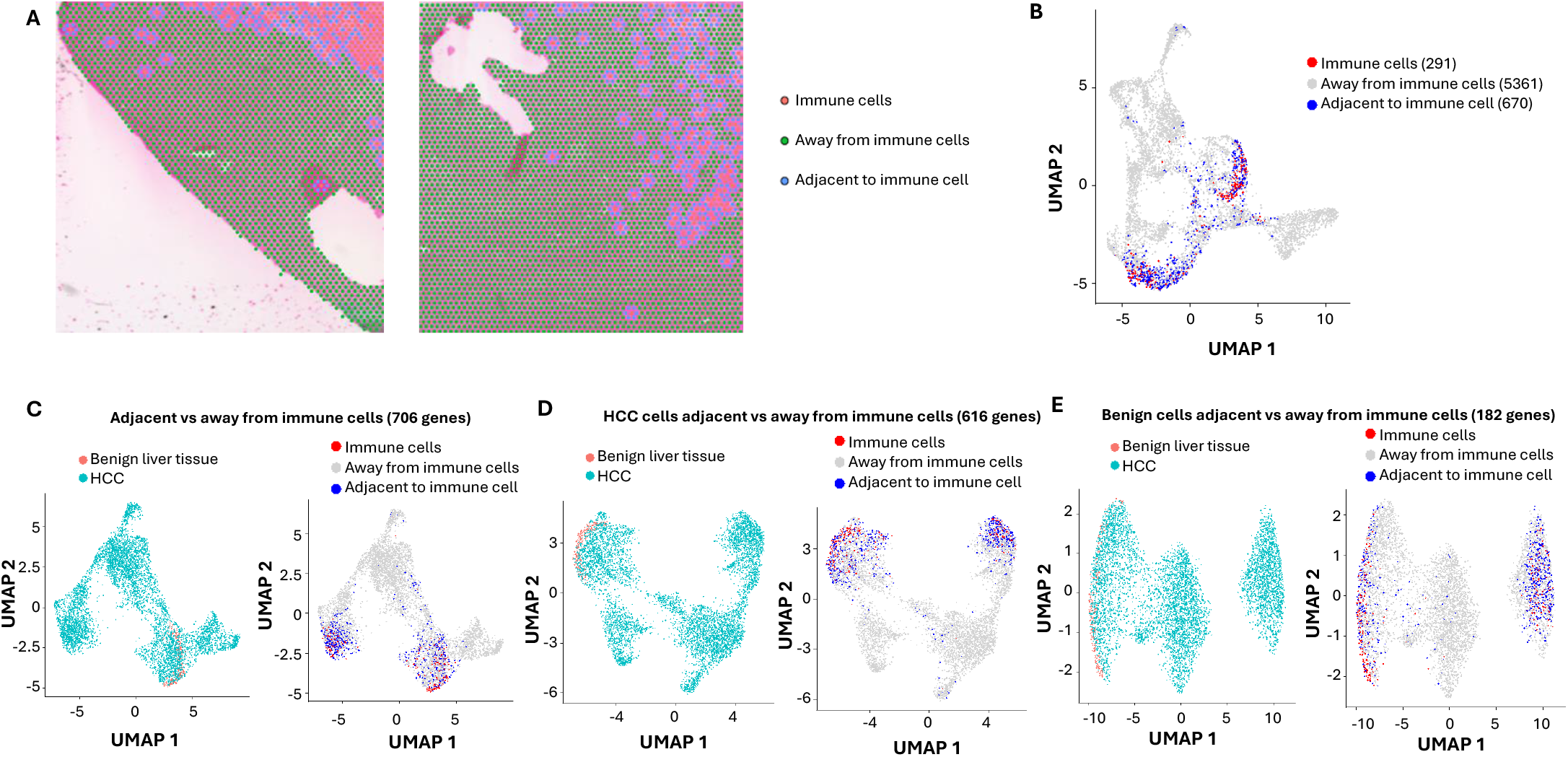
Impact of Immune cells on HCC and benign liver cells with ultra-depth sequencing. (A) Spatial visualization of immune cell-enriched, non-immune HCC, and benign liver cell spots. Immune cell-enriched spots are indicated in pink, while cells immediately adjacent to immune spots are labeled in blue. All cells away from immune spots are labeled in green. (B) The distributions of immune cell-enriched spots and cells adjacent to immune spots in UMAP clusters. (C) UMAP distributions of HCC and benign liver cells based on 706 differential expressed genes between cells adjacent to and away from immune spots (left) or UMAP distributions of immune spots, spots away from immune cells, and spots adjacent to immune cells (right). (D) UMAP distributions of HCC and benign liver cells based on 616 differential expressed genes between HCC cells adjacent to and away from immune spots (left) or UMAP distributions of immune spots, spots away from immune cells, and spots adjacent to immune cells (right). (E) UMAP distributions of HCC and benign liver cells based on 182 differential expressed genes between benign cells adjacent to and away from immune spots (left) or UMAP distributions of immune spots, spots away from immune cells, and spots adjacent to immune cells (right).

### Impact of immune microenvironment on HCC cells

To investigate the impact of immune cells on their surrounding microenvironment, differential expression analyses were performed to identify the variable genes between the spots adjacent to immune cells and the spots away from immune cells. Seven hundred and six genes were found to have differential expression between the two groups (figure 2C). When analyzing HCC cells’ responses to immune cell-enriched spots, 616 genes were found to be differentially expressed (figure 2D). While the HCC cells adjacent to immune cell-enriched spots were down on lipid metabolism, post-translational protein phosphorylation pathways, acute phase response, integrin cell surface interaction, and extracellular matrix organization pathways were up (Supplemental Table 18). For benign liver cells adjacent to immune cell-enriched spots, only 182 genes were differentially expressed versus benign liver cells away from the immune cell-enriched spots (Figure 2E). Benign liver cells adjacent to immune cell-enriched spots have down-regulation of genes in acute phase response, lipid metabolism, and post-translational protein phosphorylation pathways, while up-regulated in IL12 signaling and neutrophil extracellular trap signaling pathways (supplemental table 19), reflecting responses to cytokine release from the immune cells. Interestingly, the differences in response to immune cells between HCC and benign liver lay in the dramatic down-regulation of gene expressions in the molecular mechanism of cancer and Rho GTPase pathways in the HCC cells (supplemental table 20). These findings suggest that the primary role of immune cells is to shut down the cancer signaling pathways in liver cancer but to spare such impact in the benign liver.

### Analysis of standard-depth sequencing

Standard-depth sequencing was performed as a control to analyze the impact of ultra-depth sequencing. The mean reads in our standard-depth sequencing were 17,914-39,237 per spot. The median numbers of genes identified per spot were 1,322 to 2,769. Using the same algorithm as described for the ultra-deep sequencing (top 3000 variable genes, supplemental table 21), 15 distinct clusters were identified (Supplemental figure 3, supplemental tables 22-36). Most of these clusters were similar in terms of pathway distributions to those identified for ultra-depth sequencing. Benign liver cells were only limited to cluster 9. When immune markers were analyzed (figure 3A-B), the standard-depth produced fewer immune cell-enriched spots than the ultra-depth (271 vs 289). Only 633 genes were found differentially expressed between cells adjacent to and away from the immune cell-enriched spots (versus 706 genes for ultra-depth, figure 3C). When HCC adjacent to immune cell-enriched spots were analyzed in comparison to HCC cells away from the immune cell-enriched spots, standard-depth sequencing showed fewer genes (582 vs 616, figure 3D) and pathway affected (Supplemental table 37, 420 versus 462). In addition, standard-depth sequencing also showed fewer genes (96 versus 182, figure 3E) and pathways (Supplemental Table 38, 84 versus 137) impacted by the immune cells in the benign liver tissues. Some of the pathways showed opposite directions between standard-depth and ultra-depth sequencing. These results suggest that ultra-depth sequencing may significantly improve the analysis.

**Figure 3.**
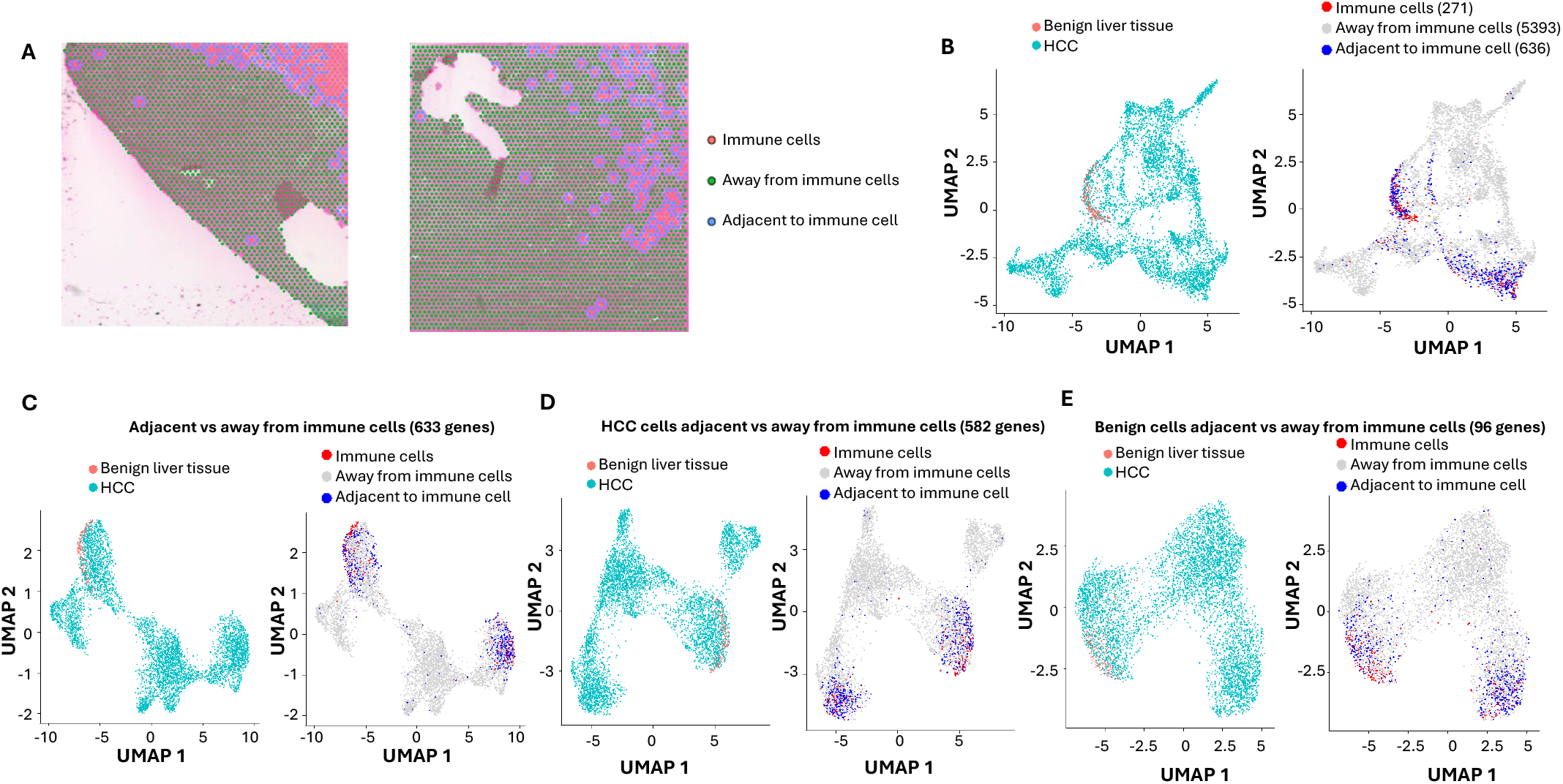
Impact of Immune cells on HCC and benign liver cells with standard-depth sequencing. (A) Spatial visualization of immune cell-enriched, non-immune HCC, and benign liver cell spots. Immune cell-enriched spots are indicated in pink, while cells immediately adjacent to immune spots are labeled in blue. All cells away from immune spots are labeled in green. (B) The distributions of immune cell-enriched spots and cells adjacent to immune spots in UMAP clusters. (C) UMAP distributions of HCC and benign liver cells based on 633 differential expressed genes between cells adjacent to and away from immune spots (left) or UMAP distributions of immune spots, spots away from immune cells, and spots adjacent to immune cells (right). (D) UMAP distributions of HCC and benign liver cells based on 582 differential expressed genes between HCC cells adjacent to and away from immune spots (left) or UMAP distributions of immune spots, spots away from immune cells, and spots adjacent to immune cells (right). (E) UMAP distributions of HCC and benign liver cells based on 96 differential expressed genes between benign cells adjacent to and away from immune spots (left) or UMAP distributions of immune spots, spots away from immune cells, and spots adjacent to immune cells (right).

To investigate what differential genes were identified between these two approaches, the genes from standard-depth were matched with ultra-depth sequencing. As shown in Figure 4, 75% (459/616) genes of ultra-depth sequencing from HCC regions impacted by immune cell-enriched spots were also identified by standard-depth. Surprisingly, 123 genes identified through standard-depth were not found in ultra-depth. For benign liver regions, only 34% (61/182) of genes of ultra-depth were found in the standard-depth sequencing. When the differences between HCC and benign liver immune-impacted genes were analyzed, only 43% of ultra-depth genes were found in the standard-depth. These results indicate that ultra-depth sequencing uncovered large numbers of biologically significant genes and pathways that standard-depth sequencing did not find.

**Figure 4.**
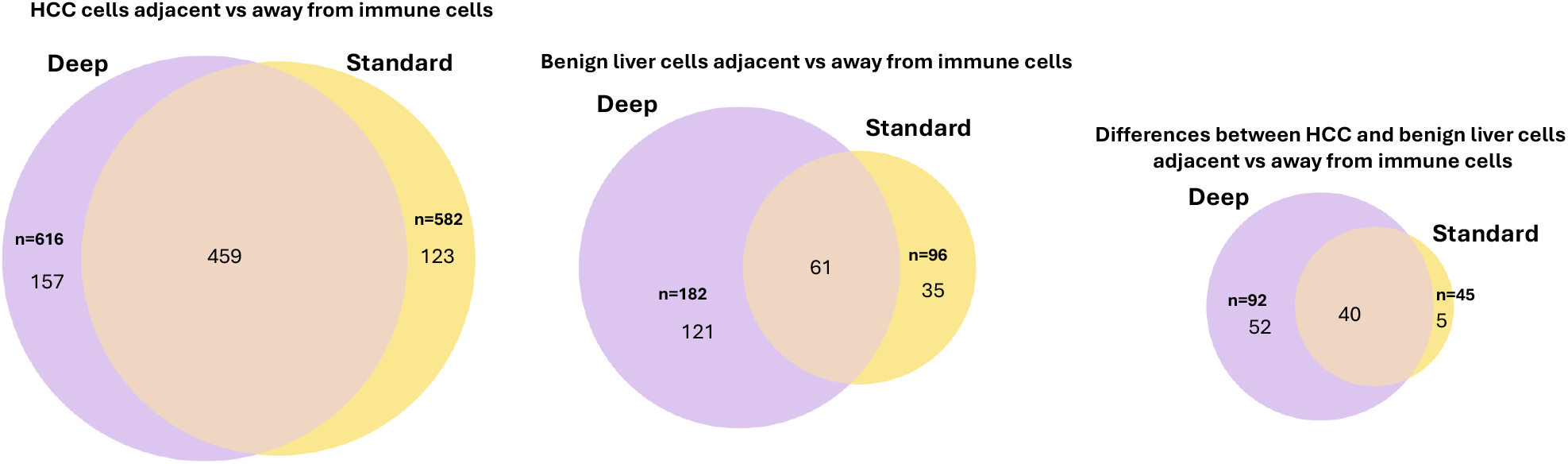
Impact of ultra-depth sequencing on differentially expressed gene discovery. Left: Venn diagram of differentially expressed genes between HCC cells adjacent to versus away from immune cells with ultra-depth or standard-depth sequencing; Middle: Venn diagram of differentially expressed genes between benign liver cells adjacent to versus away from immune cells with ultra-depth or standard-depth sequencing; Left: Venn diagram of the differences of differentially expressed genes between HCC and benign liver cells adjacent vs away from immune cells.

## Discussion

HCC is highly heterogeneous. The genotype of HCC may vary from region to region. Our study confirmed that large variations in gene expressions occurred in different regions of the cancer. Such variations were quite distinct from each other, suggesting that subclones of cancer cells had significant evolution from their origin. Underlying these gene expression variations were probably new genome mutations or chromosomal rearrangements. Indeed, recent long-read single-cell sequencing suggests that extensive mutation evolutions occurred in a small region of HCC^8^. The mutation evolution may drive the gene expression alterations that produce the cancer phenotype^27^. The exact mechanisms that induce the genetic mutations remain elucidated. However, it is likely that the DNA repair mechanism in HCC is defective. Such a defect may lead to a cascade of mutation accumulation and changes in gene expression patterns.

The tumor microenvironment has long been known to impact cancer development. Our study showed significant immune cells infiltrating both the cancer and benign liver areas. A large number of genes showed distinct responses to the presence of these immune cells. There were significant differences in response to immune cells between benign liver and HCC. There were 3-fold more genes and pathways altered in HCC in comparison with benign liver cells in response to immune cells, even though the immune cells were less abundant in the immune cell-enriched spots of the HCC area. One of the qualitative differences between the HCC and benign liver responses to immune cells is the genes of the acute response phase pathway: Genes such as SOD2 or ORM1 were up-regulated in HCC but down-regulated in benign liver. These differential responses to immune cells may suggest an innate difference in the mechanism of cytokine signaling between cancer and benign liver cells. These differences may result from the differences in the immune/somatic cell interaction or the differences in cellular sensitivity to cytokines secreted by the lymphocytes. The lack of vigorous responses from the benign liver tissues can be interpreted as normal immune adaptation.

Ultra-depth sequencing appears to offer significant advantages in identifying differentially expressed genes and pathways, particularly if the gene expression levels are not very high. One interesting finding is that ultra-depth sequencing did not cover all the differentially expressed genes discovered by 10-fold lower-depth sequencing. This suggests that even ultra-depth sequencing does not escape significant sampling errors. However, the sampling error rate could be higher in standard-depth sequencing. Neither ultra-depth nor standard-depth sequencing eliminates false negative discoveries. However, ultra-depth sequencing produced significantly more differentially expressed genes and thus uncovered more mechanisms that are important to understanding the spatial relationship and interaction between immune and cancer cells, or between cancerous and benign cells. As we reach the era of ultra-affordable sequencing, ultra-depth spatial sequencing may present an important opportunity to decipher the mechanisms of cancer development.

## Acknowledgment

This work is in part supported by grants from the National Cancer Institute (1R56CA229262-01 to JHL), the National Institute of Digestive Diseases and Kidney (P30-DK120531-01, YPY, JHL, SL), the National Institutes of Health (UL1TR001857 and S10OD028483 to SL), Innovation in Cancer Informatics (SL), and The University of Pittsburgh Clinical and Translational Science Institute (JHL).

## Conflict of interest

SL, YPY, BGR, JJL, and JHL declare no conflict of interest, while CO, TBY, MK, and KM are employees of Element Biosciences, Inc.

## Contribution

JHL, YPY, SL, TBY and CO conceived the idea. YPY, BGR, TBY, CO, and JHL contributed the experiment design. BGR, CO, TBY, MK, and KM performed the experiments. SL, CO, YPY, BJK, JJL, and JHL performed the analyses.

---

**Supplemental Figure 1. Spatial visualization of immune cell composition in slides.** HCC and benign liver areas are indicated. Each cell type is indicated by the colorization of a miniature pie chart. Top panel: Ultra-depth sequencing; Bottom: Standard-depth sequencing.

**Supplemental Figure 2. Immune cell composition is immune cell-enriched spots of HCC and benign liver.** Average fractions of each type of immune cells of the immune cell-enriched spots were shown. Left: Ultra-depth sequencing; Right: Standard-depth sequencing.

**Supplemental Figure 3. Spatial distribution of HCC and benign liver clusters with standard-depth sequencing.** (A) Spatial visualization of cell clusters in slides. Each cluster distribution is indicated by its unique color. (B) UMAP distributions of 15 spatial clusters.

## Notes

https://www.ncbi.nlm.nih.gov/geo/query/acc.cgi?acc=GSE283406

## References

1 Jemal, A. et al. Global cancer statistics. CA: a cancer journal for clinicians 61, 69–90 (2011).

2 Jemal, A. et al. Global cancer statistics. CA: a cancer journal for clinicians (2012).

3 McGuire, S. World Cancer Report 2014. Geneva, Switzerland: World Health Organization, International Agency for Research on Cancer, WHO Press, 2015. Adv Nutr 7, 418–419, doi:10.3945/an.116.012211 (2016).

4 Siegel, R. L., Miller, K. D., Fuchs, H. E. & Jemal, A. Cancer statistics, 2022. CA: a cancer journal for clinicians 72, 7–33, doi:10.3322/caac.21708 (2022).

5 Giaquinto, A. N. et al. Cancer statistics for African American/Black People 2022. CA: a cancer journal for clinicians, doi:10.3322/caac.21718 (2022).

6 Khemlina, G., Ikeda, S. & Kurzrock, R. The biology of Hepatocellular carcinoma: implications for genomic and immune therapies. Mol Cancer 16, 149, doi:10.1186/s12943-017-0712-x (2017).

7 Yu, Y. P., Liu, S., Geller, D. & Luo, J. H. Serum Fusion Transcripts to Assess the Risk of Hepatocellular Carcinoma and the Impact of Cancer Treatment through Machine Learning. The American journal of pathology 194, 1262–1271, doi:10.1016/j.ajpath.2024.02.017 (2024).

8 Liu, S. et al. Long-read single-cell sequencing reveals expressions of hypermutation clusters of isoforms in human liver cancer cells. eLife 12, doi:10.7554/eLife.87607 (2024).

9 Kader, M., Yu, Y. P., Liu, S. & Luo, J. H. Immuno-targeting the ectopic phosphorylation sites of PDGFRA generated by MAN2A1-FER fusion in HCC. Hepatol Commun 8, doi:10.1097/HC9.0000000000000511 (2024).

10 Kader, M. et al. Therapeutic targeting at genome mutations of liver cancer by the insertion of HSV1 thymidine kinase through Cas9-mediated editing. Hepatol Commun 8, doi:10.1097/HC9.0000000000000412 (2024).

11 Zuo, Z. H. et al. Oncogenic Activity of Solute Carrier Family 45 Member 2 and Alpha-Methylacyl-Coenzyme A Racemase Gene Fusion Is Mediated by Mitogen-Activated Protein Kinase. Hepatol Commun 6, 209–222, doi:10.1002/hep4.1724 (2022).

12 Liu, S. et al. Transcriptome and Exome Analyses of Hepatocellular Carcinoma Reveal Patterns to Predict Cancer Recurrence in Liver Transplant Patients. Hepatol Commun 6, 710–727, doi:10.1002/hep4.1846 (2022).

13 Luo, J. H. et al. Pten-NOLC1 fusion promotes cancers involving MET and EGFR signalings. Oncogene 40, 1064–1076, doi:10.1038/s41388-020-01582-8 (2021).

14 Yu, Y. P. et al. Detection of fusion transcripts in the serum samples of patients with hepatocellular carcinoma. Oncotarget 10, 3352–3360 (2019).

15 He, D. M. et al. Oncogenic activity of amplified miniature chromosome maintenance 8 in human malignancies. Oncogene 36, 3629–3639, doi:10.1038/onc.2017.123 (2017).

16 Chen, Z. H. et al. Targeting genomic rearrangements in tumor cells through Cas9-mediated insertion of a suicide gene. Nature biotechnology 35, 543–550, doi:10.1038/nbt.3843 (2017).

17 Chen, Z. H. et al. MAN2A1-FER Fusion Gene Is Expressed by Human Liver and Other Tumor Types and Has Oncogenic Activity in Mice. Gastroenterology 153, 1120–1132, doi:10.1053/j.gastro.2016.12.036 (2017).

18 Nalesnik, M. A. et al. Gene deletions and amplifications in human hepatocellular carcinomas: correlation with hepatocyte growth regulation. The American journal of pathology 180, 1495–1508 (2012).

19 Luo, J. H. et al. Transcriptomic and genomic analysis of human hepatocellular carcinomas and hepatoblastomas. Hepatology (Baltimore, Md 44, 1012–1024 (2006).

20 Li, Q., Zhang, X. & Ke, R. Spatial Transcriptomics for Tumor Heterogeneity Analysis. Front Genet 13, 906158, doi:10.3389/fgene.2022.906158 (2022).

21 Arora, R. et al. Spatial transcriptomics reveals distinct and conserved tumor core and edge architectures that predict survival and targeted therapy response. Nature communications 14, 5029, doi:10.1038/s41467-023-40271-4 (2023).

22 Liu, S. et al. Utility analyses of AVITI sequencing chemistry. BMC genomics 25, 778, doi:10.1186/s12864-024-10686-4 (2024).

23 Yu, Y. P. et al. Novel fusion transcripts associate with progressive prostate cancer. The American journal of pathology 184, 2840–2849 (2014).

24 Hao, Y. et al. Dictionary learning for integrative, multimodal and scalable single-cell analysis. Nature biotechnology 42, 293–304, doi:10.1038/s41587-023-01767-y (2024).

25 Ma, L. et al. Multiregional single-cell dissection of tumor and immune cells reveals stable lock-and-key features in liver cancer. Nature communications 13, 7533, doi:10.1038/s41467-022-35291-5 (2022).

26 Ma, Y. & Zhou, X. Spatially informed cell-type deconvolution for spatial transcriptomics. Nature biotechnology 40, 1349–1359, doi:10.1038/s41587-022-01273-7 (2022).

27 Liu, S. et al. Targeted transcriptome analysis using synthetic long read sequencing uncovers isoform reprograming in the progression of colon cancer. Commun Biol 4, 506, doi:10.1038/s42003-021-02024-1 (2021).

