## Supplemental figures 1-3 for "Deep Spatial Sequencing Revealing Differential Immune Responses in Human Hepatocellular Carcinoma"

### Slide 1
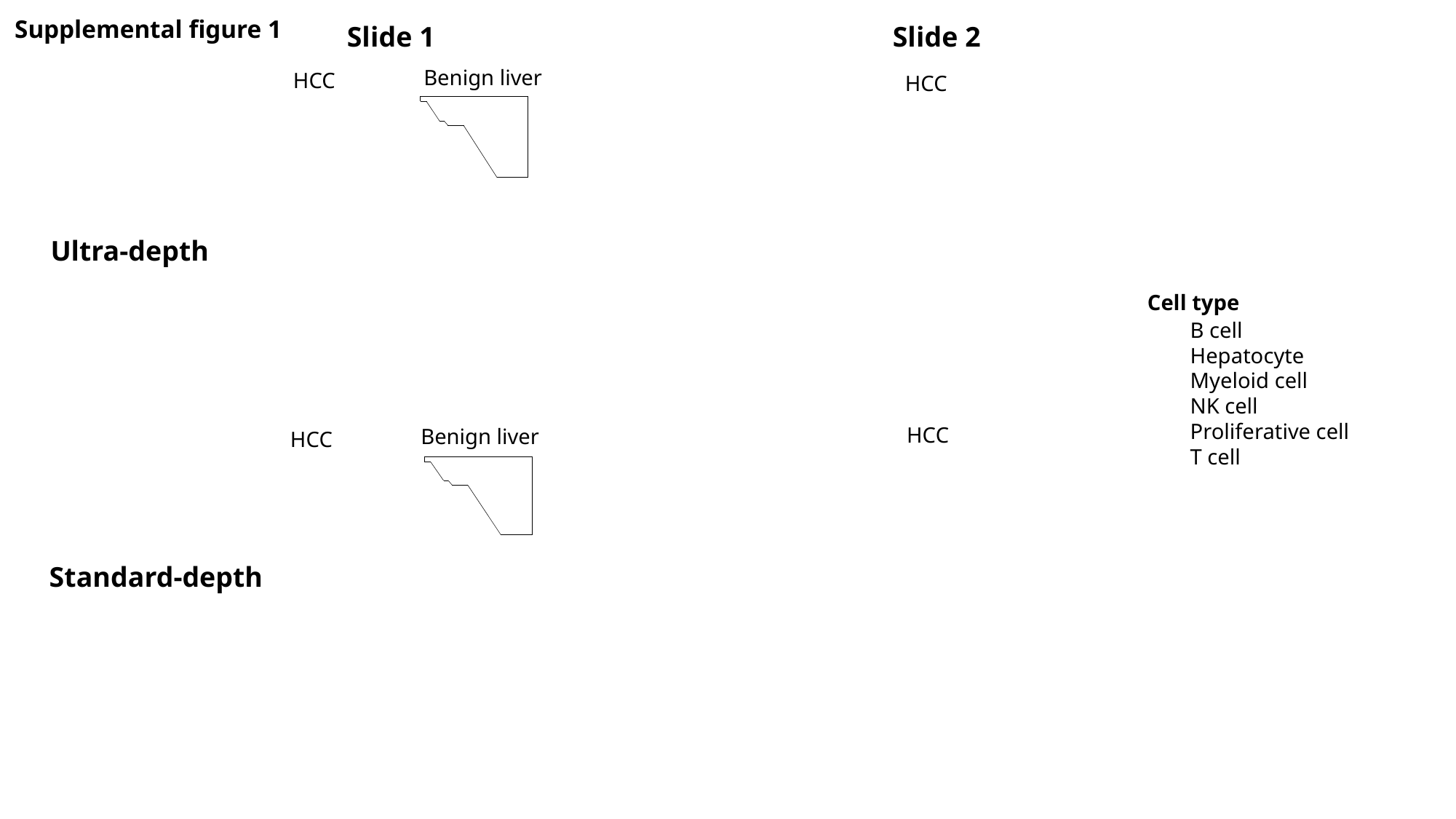

Supplemental figure 1
Slide 1					Slide 2
Benign liver
HCC
HCC
Ultra-depth
Cell type
B cell
Hepatocyte
Myeloid cell
NK cell
Proliferative cell
T cell
HCC
Benign liver
HCC
Standard-depth

### Slide 2
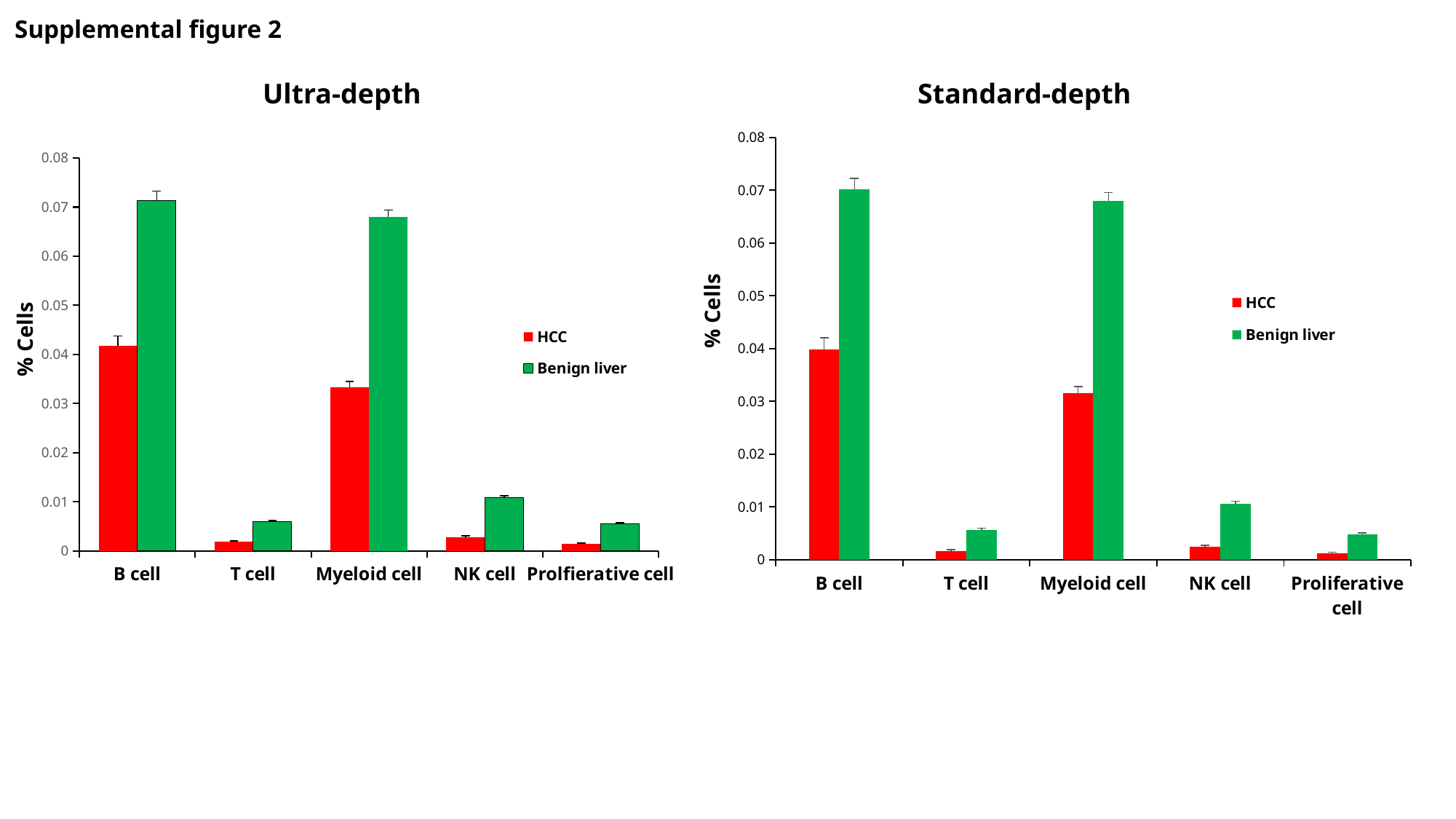

Supplemental figure 2
Ultra-depth					Standard-depth
#### Chart
| Category | HCC | Benign liver |
|---|---|---|
| B cell | 0.03987205374153369 | 0.07017623914617875 |
| T cell | 0.0016768626780730718 | 0.005688203986930585 |
| Myeloid cell | 0.03153010180149982 | 0.06794161709619145 |
| NK cell | 0.002466579539241382 | 0.01057194617761123 |
| Proliferative cell | 0.0012560982092858343 | 0.004868217050594113 |
#### Chart
| Category | HCC | Benign liver |
|---|---|---|
| B cell | 0.04173410761468632 | 0.07130338085999757 |
| T cell | 0.0018057228869019922 | 0.005893303036790473 |
| Myeloid cell | 0.033269282217222106 | 0.06797561350502426 |
| NK cell | 0.002791454059544798 | 0.010767356488762704 |
| Prolfierative cell | 0.0014081740397805746 | 0.0055293408118424064 |% Cells
% Cells

### Slide 3
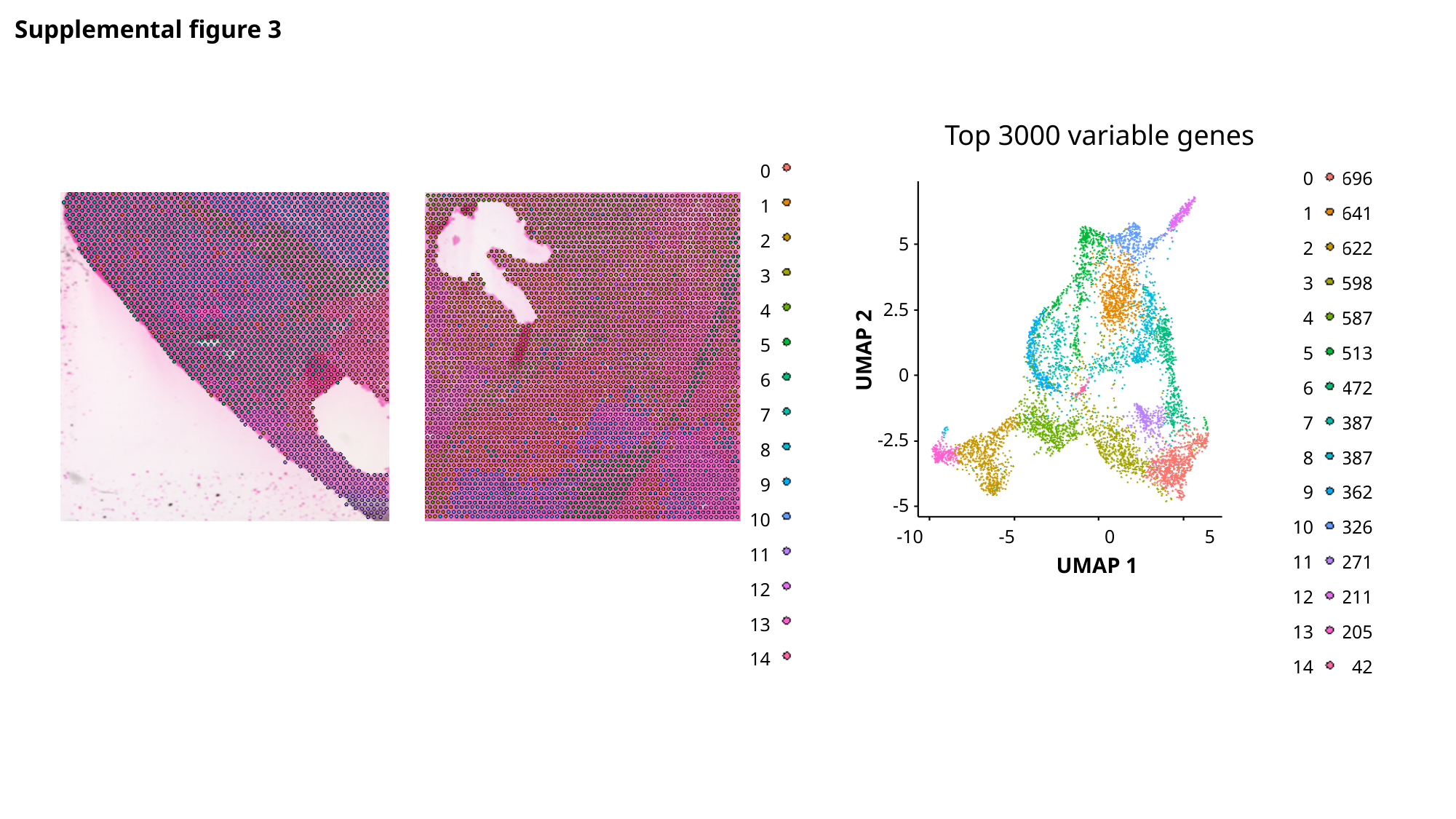

Supplemental figure 3
Top 3000 variable genes
0
1
2
3
4
5
6
7
8
9
10
11
12
13
14
0
1
2
3
4
5
6
7
8
9
10
11
12
13
14
696
641
622
598
587
513
472
387
387
362
326
271
211
205
42
5
2.5
0
-2.5
-5
UMAP 2
-10 -5 0 5
UMAP 1
